# Open source 3D printable replacement parts for the WHO insecticide susceptibility bioassay system

**DOI:** 10.1101/762849

**Authors:** Sean Tomlinson, Henrietta C. Yates, Ambrose Oruni, Harun Njoroge, David Weetman, Martin J. Donnelly, Arjen E Van’t Hof

## Abstract

**Background:** Malaria vector control and research rely heavily on monitoring mosquito populations for the development of resistance to public health insecticides. One standard method for determining susceptibility in adult mosquito populations is the World Health Organization test (WHO bioassay). The WHO bioassay kit consists of several acrylic pieces that are assembled into a unit. Parts of the kit commonly break, reducing the capacity of insectaries to carry out resistance profiling. Since there is at present only a single supplier for the test kits, replacement parts can be hard to procure in a timely fashion. Here, we present 3D printable versions for all pieces of the WHO bioassay kit.

**Results:** Using widely available polylactic acid (PLA) filament as a printing material, we were able to design and print functional replacements for each piece of the WHO bioassay kit. We note no significant difference in mortality results obtained from PLA printed tubes and WHO acrylic tubes. Additionally, we observed no degradation of PLA in response to prolonged exposure times of commonly used cleaning solutions.

**Conclusions:** Our designs can be used to produce replacement parts for the WHO bioassay kit in any facility with a 3D printer, which are becoming increasingly widespread. 3D printing technologies can affordably and rapidly address equipment shortages and be used to develop bespoke equipment in laboratories.

## Background

Malaria remains a critical public health problem across sub-Saharan Africa, with vector control — a vital part of efforts to control and eradicate malaria — relying heavily on efficacious insecticides [1]. Widespread and emerging resistance poses a significant threat to public health and is reflected by increased efforts to understand and characterize the distribution of resistant mosquito populations and associated genetic variants across endemic regions of Africa [2, 3].

The World Health Organization insecticide susceptibility test (WHO bioassay) is a standard method implemented to assess resistance in adult mosquito populations. During this test, mosquitoes are held in one of two tubes (Fig. 1a), either lined with untreated paper (control) or insecticide-impregnated paper (exposure) held in place with spring clips (Fig. 1c). Both tubes are separated by a slide unit (Fig. 1e) and slide (Fig. 1f), while the ends of the tubes are capped with a screen mesh (Fig. 1b) and screw cap (Fig. 1a). Mosquitoes are held in the insecticide tube for one hour, and the percentage mortality of exposed mosquitoes 24 hours post-exposure is a measurement for insecticide susceptibility [4]. A single experimental unit for the WHO bioassay kit is comprised of 2 mesh screens, two screw caps, two tubes, four spring clips, one slide unit, one slide (Fig. 1).

**Figure 1.**
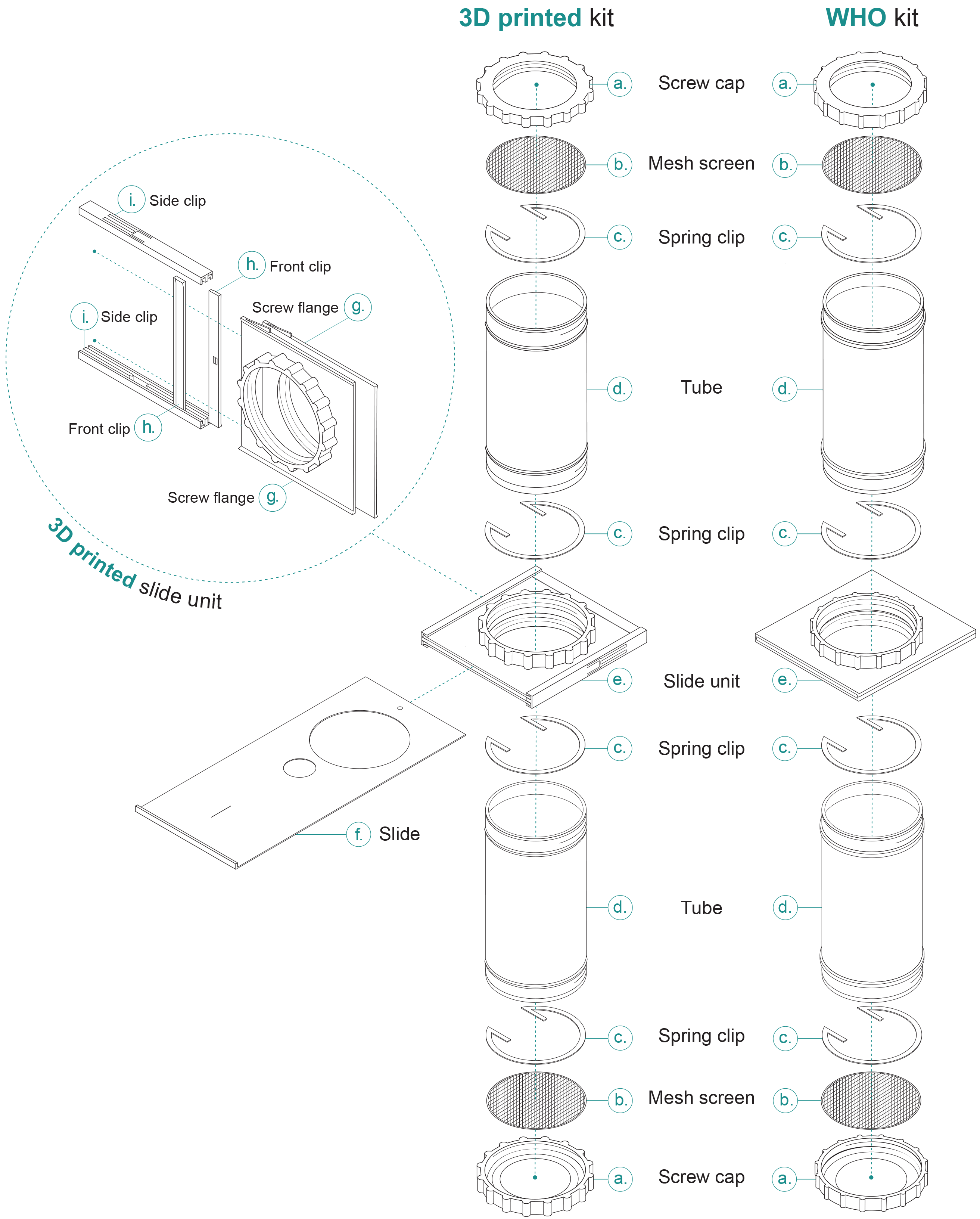
World Health Organisation bioassay kit and equivalent 3D printed parts. Cutaway shows the assembly of the 3D printed version of the slide unit. For illustration purposes, the WHO bioassay slide has been omitted but is visually identical to f.

Certain parts of the WHO bioassay kit are more liable to become worn, damaged or lost, causing a reduced capacity of insectaries to conduct bioassays. Most notably, in our experience, the mesh screen can become easily lost or damaged during cleaning; the slide unit is subject to friction from the slide and when combined with the gradual weakening of the chemical bond through repeated uses and washes, frequently splits; and springs are lost during washing procedures. Long shipping times and associated costs mean that replacing lost or damaged parts can become economically or logistically unviable. To address these problems, we used computer-aided design software to produce 3D printable versions of the parts that comprise the WHO bioassay kit.

Accurate, reliable and affordable 3D printing technologies are now commercially available. The most common 3D printer form utilizes a Cartesian axis system to control the deposition of molten plastic filament onto a print surface, in a process called fused filament fabrication (FFF). Many different plastics and materials can be used for 3D printing, such as polylactic acid (PLA), acrylonitrile butadiene styrene (ABS), nylon, polyethylene terephthalate glycol (PET-G) and polycarbonate. PLA is widely available and is suitable for use in most laboratory plastic equipment. Indeed, 3D printing technologies are increasingly being used in research settings [5]. The glass transition temperature of PLA is 60 - 65°C with a melting temperature of ~180°C, meaning in cold or low-temperature settings PLA is thermally stable.

Here, we present 3D printable replacement parts for the WHO bioassay kit which print without the need for tools or glue; and which interface with existing WHO bioassay parts. We discuss the design challenges, modifications from existing WHO bioassay kits and files needed to print replacement parts for the WHO bioassay kit.

## Methods

### Designing 3D models

We used SketchUp (Trimble Inc.) and OpenSCAD (Marius Kintel, Openscad.org) to create the 3D model files in the stereolithography (STL) format needed to enable 3D printing of parts. Some parts were technically difficult or impossible to directly replicate using current FFF 3D-printing. In these cases, we modified the existing design to allow printing, while retaining the same physical function.

Support material is plastic printed alongside the desired part to prevent necessary plastic overhangs from dropping below their intended position. This support material is printed in such a way that it is easily detached from the finished piece; however, its inclusion leads to longer print times and higher plastic consumption. Around the circumference of the tube, two rims are present to provide a positive stop for when the tubes are fully inserted into the slide unit. On the original WHO bioassay tube, these rims are squared on the edges, replicating this feature would require support material during printing. To reduce print time, plastic consumption and potential interference with tube threads, the outer geometry of the rim was changed to triangular. This geometry can be printed without any lower support while retaining the function of the original part.

The slide unit has an internal section into which the gate slides. This geometry is complex; indeed, the original part is manufactured in two halves and chemically bonded together. The concept for this project required that the entire system be 3D printable, to increase accessibility and use. To be practically printable, this part needed adapting for 3D printing. Like the WHO bioassay slide unit, we created two halves and developed a method of bonding the pieces together. We designed a sliding clip method of joining two screw flanges of the slide unit. Two halves of the slide unit are printed with the addition of arrow-like notches on each side; these interface with a sliding lock clip that mechanically locks the two halves together and creates a gap for the gate to slide through (Fig. 1g, h, i).

On the inside of the slide unit are two friction nodules (Fig. 1h) that retain the slide in either the closed or open position, preventing the slide from falling out of the slide unit during handling. To address this, we designed the whole slide unit to include front clips that retain the friction nodules. These changes now necessitate some assembly of the slide unit once printed. However, the slide unit has been designed to allow hand-assembly without the need for tools. Despite the changes to this part of the WHO bioassay kit and the increase in physical size, the mechanical function remains the same.

The mesh screen used at the end of the tubes is manufactured from a flexible material that allows it to have no border. In our prototyping, we found that printed mesh screens were too weak to be handled when printed without a border. Therefore, a 3mm border was added to the CAD version of the mesh – this does not extend past the lip of the screw cap, retaining the same function as the original.

### 3D Printing

3D printing was carried out on an Original Prusa i3 MK3 and an MK3S (Prusa Research) modified with a BuildTak print surface (https://www.buildtak.eu/), using white 1.75mm PLA filament (ZIRO3D). Designed CAD models were exported as STL files. STL files must be converted to machine instructions following the G-code standard to be processed by 3D printers. This conversion process is called slicing. The STL model files were sliced using Cura 3.3.1 (Ultimaker) with the following key slicer settings: 100% infill, two shells/perimeters.

Reliably and efficiently 3D printing transparent objects is technically difficult with commercially-available 3D printers and typically results in a cloudy translucent finish. During prototyping, we identified that bright white filament – though not transparent – provides enough contrast for mosquitoes to be easily counted while viewing through the mesh screen. Commercially available WHO bioassay tubes use a green and red dot to denote both the holding and exposure side the bioassay kit, respectively. We used a permanent marker to label the corresponding printed parts with an ‘E’ (exposure) and ‘H’ (holding) (Fig. 2).

**Figure 2.**
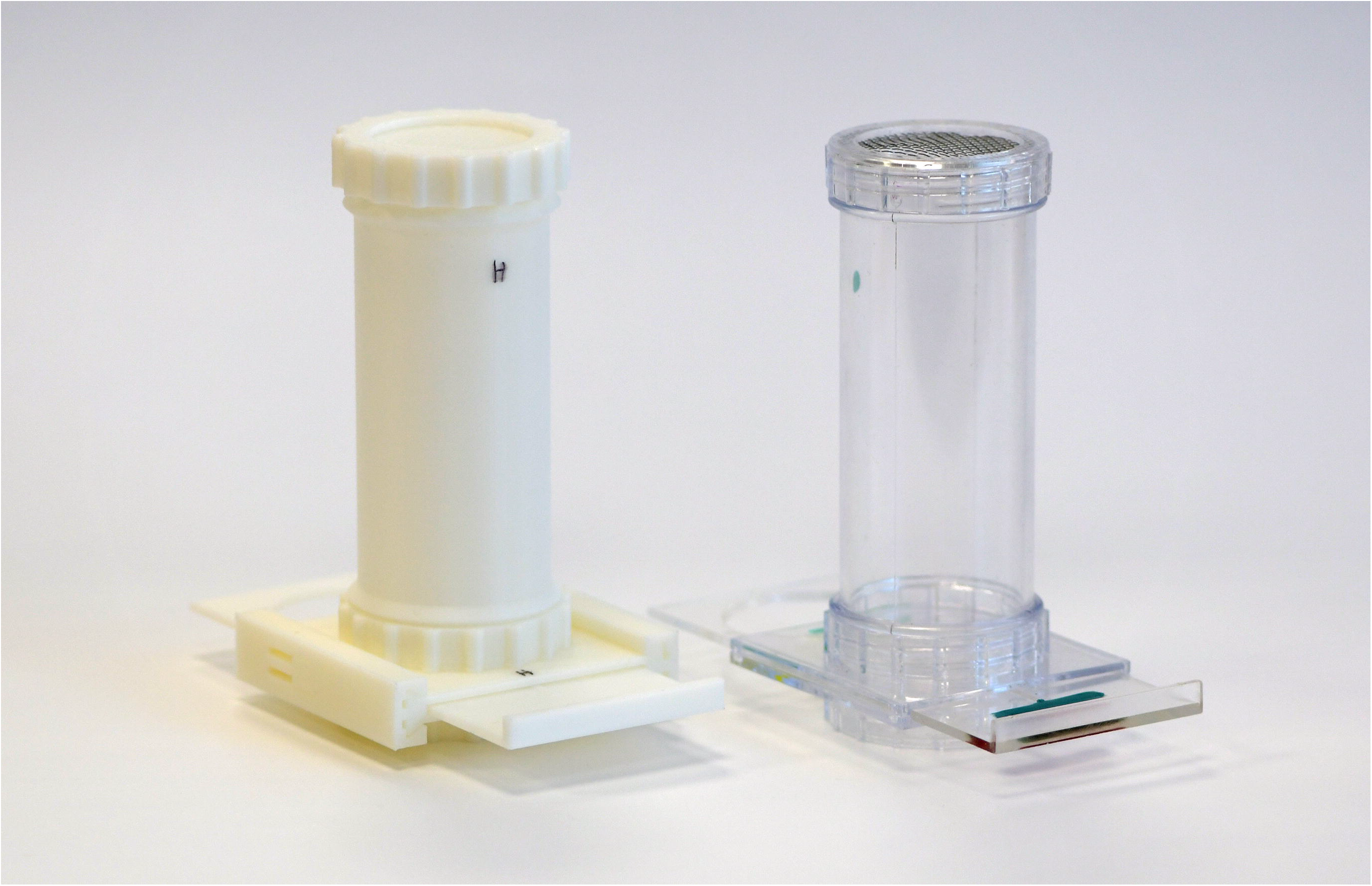
Photograph of WHO bioassay component and their corresponding 3D printed versions. E and H are used to denote the exposure and holding sides, respectively, whereas on the WHO kit a red and green dot is used.

### Bioassay testing

To ensure that the 3D printed tubes performed in a similar manner to the acrylic tubes we performed susceptibility testing using standard 4% WHO diagnostic dose of dichlorodiphenyltrichloroethane (DDT) insecticide on two Kenyan laboratory strains of *Anopheles gambiae s.s.* (Kilifi and Mbita). Batches of ~25 3 – 5 day old female mosquitoes, were exposed in each tube. The number of replicate exposures was as follows: Kilifi WHO tubes n=10, Kilifi 3D tubes n=11, Mbita WHO tubes n=7, Mbita 3D tubes n=7. Percentage mortality was recorded after the 24-hour holding period. All mosquito rearing was conducted at the Liverpool School of Tropical Medicine insectaries, following standard operating procedures. The Mbita strain was collected at Mbita Point, Kenya in 1999, and has been maintained as a laboratory strain since this time. The Kilifi strain was collected in the Kilifi County, Kenya in 2012. The colony is maintained by both the Liverpool School of Tropical Medicine and Kenya Medical Research Institute.

### PLA reactivity with bioassay solutions

To assess whether the PLA would interact with solutions that are commonly used during the bioassay protocol, we exposed PLA parts to 4 different solutions to observe any degradation of the plastic. (1) Cotton pads soaked with 10% sucrose solution, typically used to feed mosquitoes during the recovery period, were placed on six mesh screens for seven days. Cotton pads were soaked daily with fresh 10% sucrose solution to replace evaporated solution. (2) Four slides were submerged in 3% Rely+On Virkon (Lanxess) for five days. (3) Four slides were submerged in 5% Decon 90 (Decon Laboratories Ltd.) for five days. (4) Six screw caps were submerged in 70% ethanol for five days.

## Results

### 3D printing

Designed and printed parts interface as expected with current WHO bioassay parts. The printed kits assembled easily without the need for additional tools. CAD and STL files produced are available at https://github.com/SeanTomlinson30/3D-Printable-WHO-Bioassay-Parts.

### Bioassay testing

Bioassays with 4% DDT using the Mbita and Kilifi strains showed no significant difference in 24-hour mortality (Figure 3) for measurements between 3D printed and WHO bioassay kits. We also observed that mosquitoes can sugar feed through the 3D printed mesh screens.

**Figure 3.**
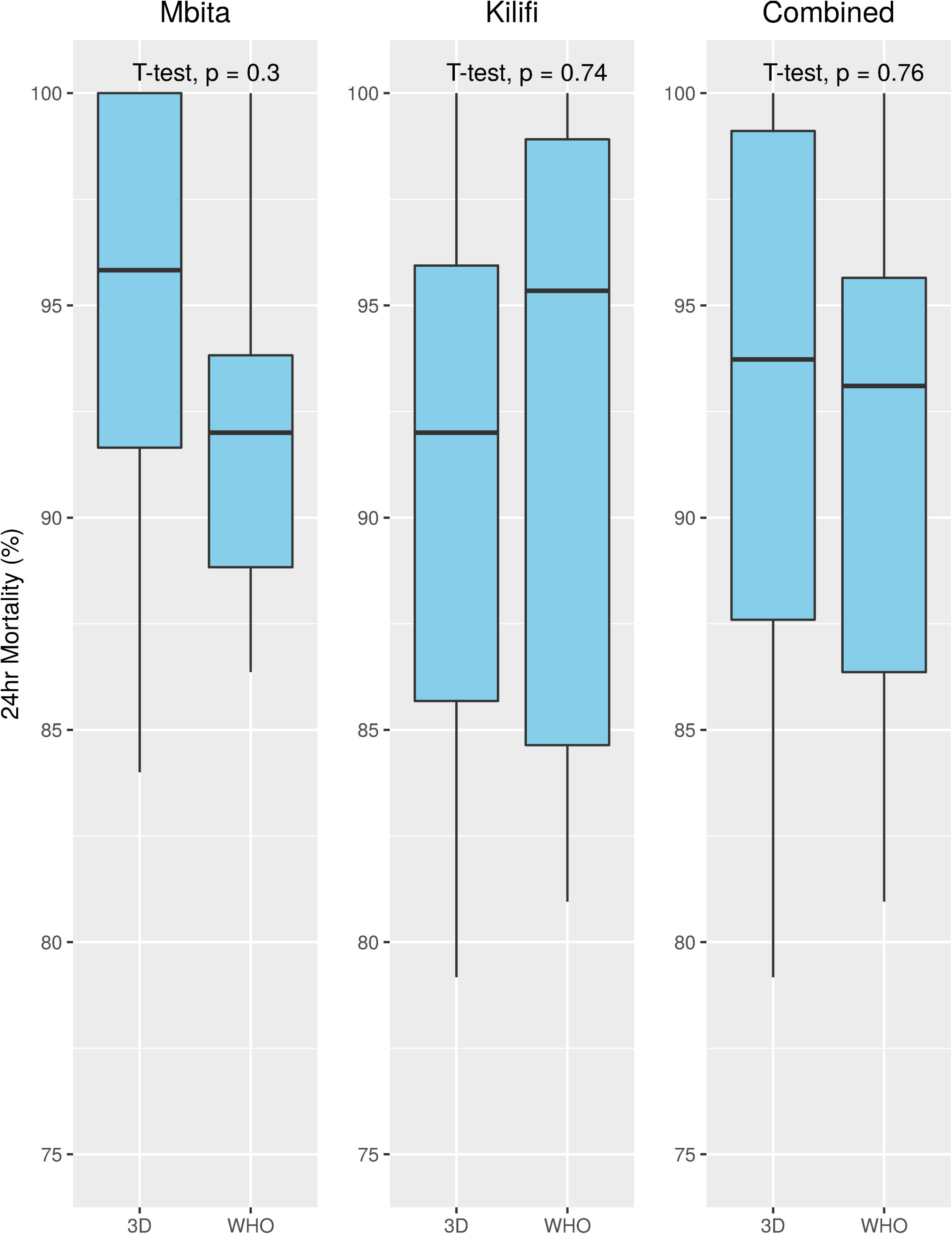
Bioassay results comparing mortality between the WHO bioassay kit and 3D printed kit for Mbita, Kilifi and their combined results. A t-test was used to compare means for each test.

### PLA Reactivity with Bioassay Solutions

After exposure to 10% sucrose, 70% ethanol, 3% Rely+On Virkon (Lanxess) and 5% Decon 90 (Decon Laboratories Ltd.), we observed no signs of degradation of the PLA strength, tensibility, surface colour or size.

## Discussion

We have developed, and provide here, printable versions of all pieces that compose a WHO bioassay kit. We see the primary use case for these parts as a replacement library for missing and damaged parts of an original WHO bioassay kit. Bioassay data for DDT exposure indicate no significant difference between 3D printed and WHO bioassay kits; although, other insecticides/strain combinations may react differently when interacting with 3D printed materials. Anecdotally, in our insectaries, we find that the most in-demand 3D printed replacement parts are the slide unit and mesh screen, with tubes being the most durable parts and least likely to be needed.

The design challenges of 3D printing the WHO bioassay kit necessitated some changes to the geometry of individual parts. Most notably, to retain all functionality, the 3D printable slide unit had to be printed as six individual pieces that are assembled. In addition to showing no functional differences during operation and manual handling, because the 3D printed slide unit does not use chemical bonding, it is more durable to general wear and less likely to become damaged, in terms of splitting. Though, we do note that when using PLA as a 3D printing material, operators must be cognizant of the effect of hot temperatures causing material deformation.

## Conclusions

We present files that allow printing of all parts of the WHO bioassay kit. To achieve this, we replicated existing parts in CAD software, modifying and adapting the designs where necessary to permit 3D printing. The printed parts interface with standard WHO bioassay kits and in the case of full 3D printed kits, produce results not significantly different from standard WHO bioassay kits. 3D printing in laboratory environments has become more achievable thanks to the continued reduction in costs and developments in 3D printing technologies. Through the distribution of the 3D printable laboratory equipment, researchers can maintain testing capacity, reduce costs and adapt apparatus for bespoke purposes.

## List of abbreviations

ABS: acrylonitrile butadiene styrene
CAD: computer-aided design
DDT: dichlorodiphenyltrichloroethane
FFF: fused filament fabrication
PET-G: polyethylene terephthalate glycol
PLA: polylactic acid
STL: stereolithography
WHO: World Health Organisation

## Declarations

### Ethics approval and consent to participate

Not applicable.

### Consent for publication

Not applicable.

### Availability of data and material

The CAD and STL files produced are in the supplementary materials and are available at https://github.com/SeanTomlinson30/3D-Printable-WHO-Bioassay-Parts. Bioassay data and analyses scripts are found in the supplementary materials (S1, S2).

### Authors’ contributions

ST, AVH and MJD designed the study. ST and AVH designed the CAD models and 3D printed the parts. DW, HN and AO helped design the assays. AO carried out the assays. HCY performed initial prototype testing. ST wrote the manuscript. All authors read and approved the final manuscript.

## Acknowledgments

We are grateful to Manuela Bernardi, who illustrated figure 1. We also would like to thank Giorgio Praulins for providing example WHO bioassay tubes for the CAD modelling process.

## Figures and additional files

S1. Bioassay mortality data for WHO vs 3D printed kits with both Mbita and Kilifi stains. (.csv)

S2. Figure and statistics generation script (.R)

